# An Open-Source Code To Analyze Mitochondrial Intracellular Distribution From Fluorescence Microscopy Images

**DOI:** 10.64898/2026.01.06.697902

**Authors:** Giovanna C. Cavalcante, Alicia J. Kowaltowski

**Affiliations:** Departamento de Bioquímica, Instituto de Química, and Centro de Metabolismo CoMeta, Hospital Universitário, Universidade de São Paulo São Paulo, Brazil

**Keywords:** mitochondria, mitochondrial localization, mitochondrial morphology, MitoTracker, immunofluorescence

## Abstract

Mitochondria have a plethora of roles in cells, many of which are related to dynamic changes in their size, shape, and intracellular location. Mitochondrial morphology is commonly assessed by microscopy with targeted fluorescent probes. However, tools to easily estimate mitochondrial localization within a cell are still lacking. A code was designed to estimate per-cell mitochondrial radial localization (perinuclear or peripheral) from fluorescence microscopy files in a variety of formats and using different mitochondrial markers (https://github.com/cavalcantegc/mito_localization.git). Three case studies with different cell types and stainings demonstrate that mitochondrial localization can be easily extracted and plotted with this code.

## 1. Introduction

Mitochondria are cytoplasmic organelles crucial in energy metabolism and multiple other intracellular processes as diverse as controlling intracellular calcium, determining cell fate in differentiation, and maintaining redox balance. Thus, it is not surprising that mitochondrial dysfunctions are strongly linked to the development and progression of complex diseases, including metabolic disorders, neurodegenerative and cardiovascular diseases (Vercesi et al., 2018).

Among the myriad of mitochondrial characteristics that may influence function and behavior, mitochondrial morphology has attracted particular attention (Monzel et al., 2023). Morphology is a characteristic directly associated with mitochondrial dynamics, as abnormalities in fission and fusion processes can impact mitochondrial size and shape, as well as downstream positioning within the cell, influencing mitochondrial behavior (Solomon et al., 2022). For example, in brown adipose tissue, peri-lipid droplet mitochondria have more elongated morphology, low fission-fusion dynamics, low beta oxidation capacity and high ATP synthesis compared to cytoplasmic mitochondria (Benador et al., 2018). Additionally, mitochondria located in the soma, dendrites or axons of neurons display differences in function and morphology (Seager et al., 2020).

Recently, an association between mitochondrial localization in a cell and a variety of psychoses (bipolar disorder, schizophrenia, and schizoaffective disorder) has been uncovered (Haghighi et al., 2025). In that study, a set of different fluorescence-based images (nuclei, cytoskeleton, and mitochondria) were processed using novel CellProfiler pipelines to obtain mitochondrial localization. Interestingly, mitochondrial localization seems to be distinct also in skin fibroblasts from patients with psychoses versus controls: mitochondria were found to be more internally localized, farther from the cell boundary, in the psychiatric patients. This reinforces the importance of mitochondrial cellular localization as a feature associated with functional outcomes.

To the best of our knowledge, there are no tools currently available to easily estimate mitochondrial localization in a cell using only typical fluorescence images. Here, we present an open-source Python code that estimates per-cell perinuclear and peripheral mitochondria from a variety of microscopy image types.

## 2. Materials and Methods

### 2.1. Code design

A code was designed to estimate per-cell mitochondrial radial localization (perinuclear or peripheral) from fluorescence microscopy assays with mitochondrial staining, using Python 3.9.7 programming language in the PyCharm Community Edition (v. 2021.3.2) environment. The full code is openly available at https://github.com/cavalcantegc/mito_localization.git. Figure 1 shows a step-by-step workflow for code usage.

**Figure 1.**
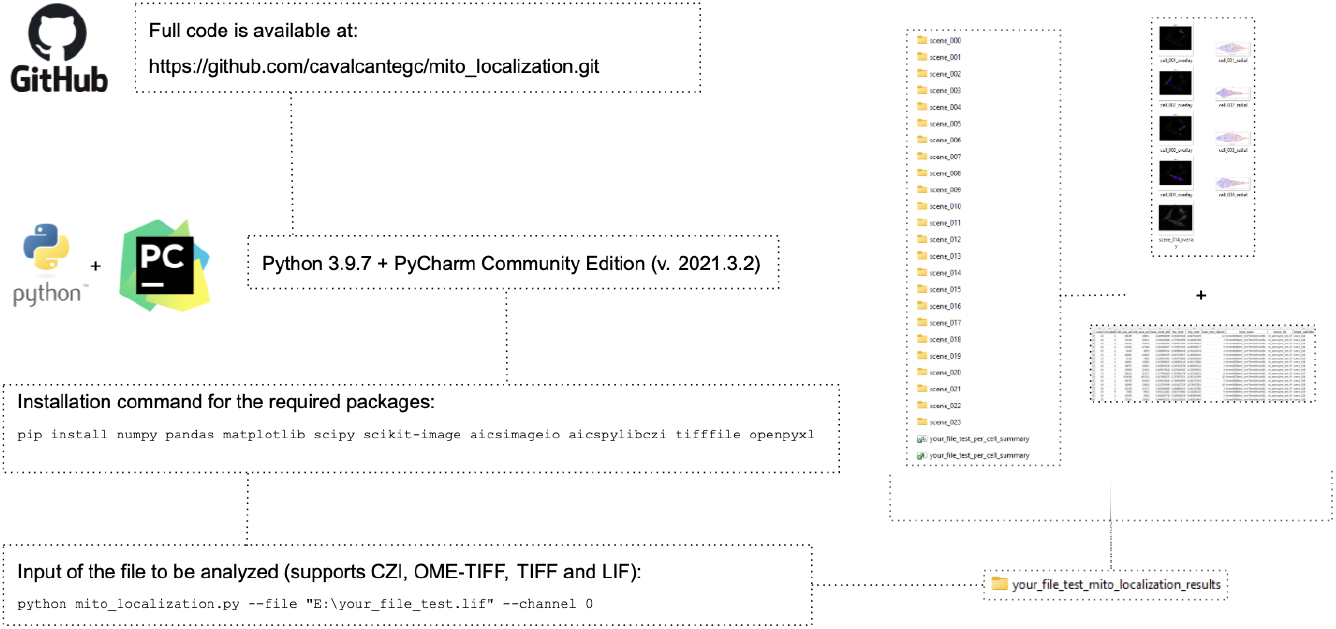
Workflow of the basic steps for code usage. The simplified input system involves adding the location of the file in the PyCharm terminal, after which the user can run the designed code with a single command line after installing the required Python packages. More instructions are given throughout the code. User-friendly outputs include a folder with individual per-cell data of each image and a summary spreadsheet.

The script estimates cell outlines from the fluorescence signal (stainings such as MitoTracker probes), and reads one or more images (micrographs captured in the assay) in each input file in a single run. It supports CZI, OME-TIFF, TIFF, and LIF formats as input, and any staining for that marks individualized mitochondria. The code was designed to generate a directory with folders for each image, including PNG violin plots (one per cell) showing the radial distribution of the mitochondrial area and an overlay for each cell. It also generates a spreadsheet with the summary of per-cell metrics, including the fraction of inner (perinuclear) and outer (peripheral) mitochondria in each labeled cell, which can then be exported to other data analysis softwares. All outputs are saved in an output folder next to the input file in the directory.

### 2.2. Applications

To demonstrate the functionality of our Python code to quantify mitochondrial intracellular distribution, we first tested data acquired by our group using human astrocytes. The code was applied to analyze a *lif* file with multiple images from assays of the cells stained with MitoTracker Deep Red FM (Invitrogen, #M22426), which accumulates in the mitochondria of living cells in an inner membrane potential-dependent manner. The nucleus area was estimated from the “void” in the captured cell. Photographs were obtained using a Leica DMi8 inverted microscope with 63x zoom.

To expand the code usage to *czi* files, we also applied it to immunofluorescence micrograph files of mouse embryonic fibroblasts (MEFs) kindly donated by Scorrano group at University of Padova, Italy. Mitochondria are marked as indicated by mitochondrial protein SDHA (succinate dehydrogenase complex flavoprotein subunit A). Photographs were obtained with the Zeiss LSM900 Airyscan confocal microscope with 63x zoom.

Moreover, we also accessed publically available datasets with microscopy images from the CRBS Cell Image Library (CIL, RRID:SCR_003510). We applied the code to the *tif* file under accession number 13731, from a study by Jin et al. (2011). The image shows HeLa cells expressing mito-YFP (green), localized to mitochondria, and stained for Tom20 (white), a mitochondrial outer membrane protein, resulting in two different fluorophores marking mitochondria. Photographs were obtained with the Zeiss LSM510 Meta microscope with a 63x zoom.

## 3. Results and Discussion

A common assay to visualize mitochondrial morphology is staining with fluorescent probes such as MitoTracker, generating images that can be analyzed by open software such as ImageJ, which has add-ons capable of generating automated quantifications of mitochondrial characteristics such as sphericity and mean volume per cell. However, subcellular mitochondrial location could not be easily analyzed with tools already at hand.

Considering this gap, we developed a user-friendly Python code to estimate per-cell mitochondrial radial localization (perinuclear or peripheral) from *lif* image files with mitochondria-targeted fluorescence, such as MitoTracker. The code produces a spreadsheet with per-cell measurements and visual outputs which facilitate the visualization of mitochondrial distribution both by using a color code (Figure 2A) and by quantifying distance from the nucleus (Figure 2C). With a single command line as input, the code identified each cell (Figure 2B) and estimated the fraction of mitochondria that are located in inner and outer portions of the cell, depicting the data as figures and quantitative spreadsheets (Figure 2D). In the representative cell depicted in Figure 2, more mitochondria are perinuclear, although a wide variety of locations is observed. Mitochondrial dynamics, particularly fission, may modulate morphogenesis and mitochondrial organization within the cell in astrocytes (Salazar et al., 2025), highlighting the importance of being able to readily quantify mitochondrial distribution in a cell.

**Figure 2.**
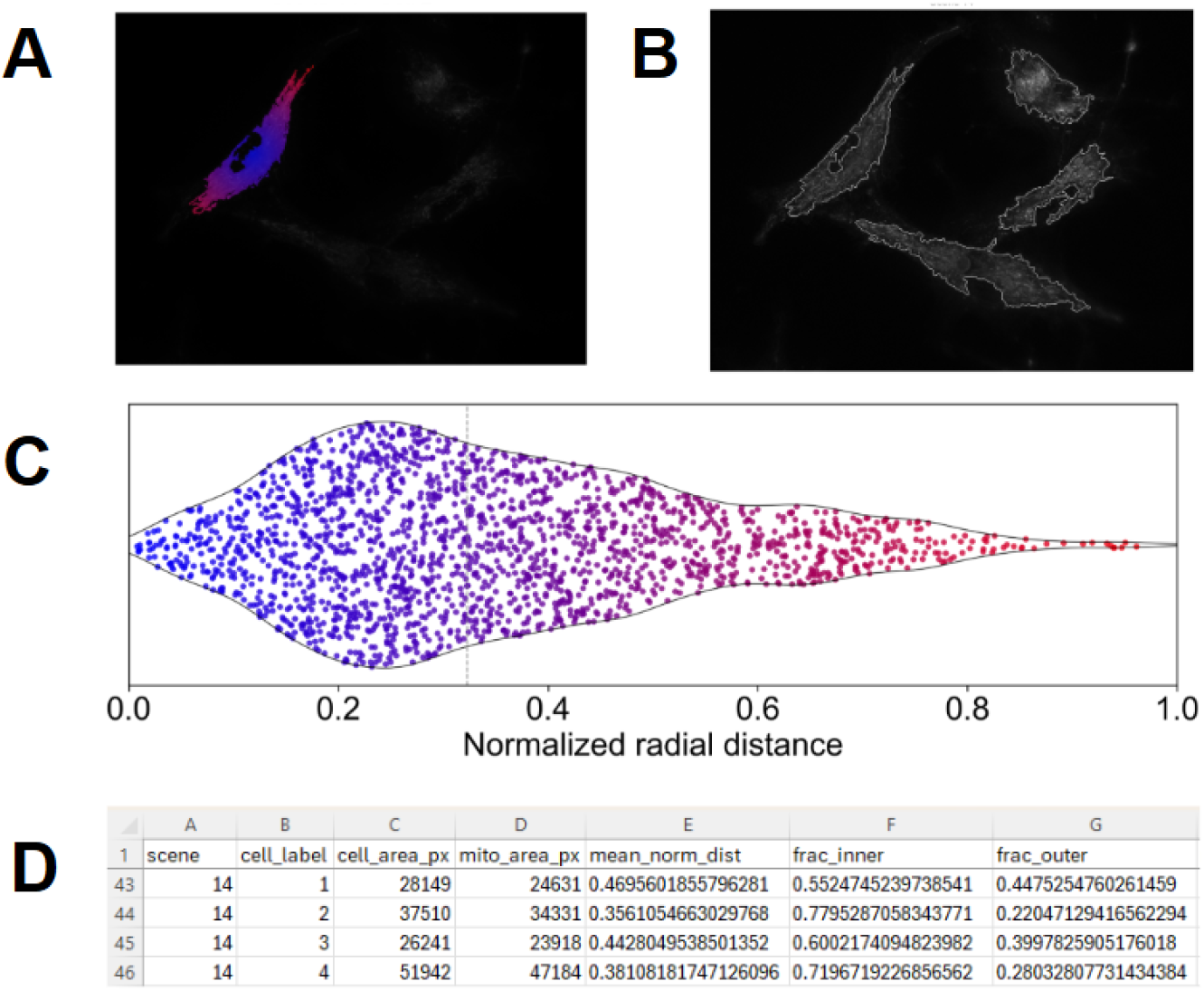
Mitochondrial localization quantification in human astrocytes. **A)** Per-cell estimates of mitochondrial localization shown a color gradient from perinuclear (blue) to peripheral (red) mitochondria obtained from analysis of a MitoTracker-stained cell image (gray). **B)** Gray layer of cells detected and mitochondria in the image. In this example, “Scene 14” shows four cells. **C)** Violin plot with the fraction of mitochondrial pixels (dots) identified from perinuclear (center, 0, in blue) to peripheral (periphery, 1, in red). **D)** Values in the spreadsheet generated for the four cells in Panel B.

In a second example, we applied the code to *czi* files in which MEF cells were stained with anti-SDHA immunofluorescence, in order to test the code for use with different mitochondrial stains and image file types. No issues were observed when changing either the file format or the fluorescence modality, and the code executed successfully. However, the test showed that when using a different cell type, adjustments were needed. MEFs are more rounded than astrocytes, with larger nucleus “voids” and sparse mitochondria forming clusters. In this image, MEFs are also touching each other, forming closer mitochondrial networks, as seen in Figure 3. Sparse mitochondrial signals can cause oversegmentation, while physically adjacent cells may be perceived as a single cell. This highlights that, as with other automated detection codes, visual checks for quality control of the cell separations and mitochondrial identification are important. Additionally, given the nature of the code is to determine the location of individual mitochondria within cells, it requires cells with at least a partially fragmented mitochondrial network.

**Figure 3.**
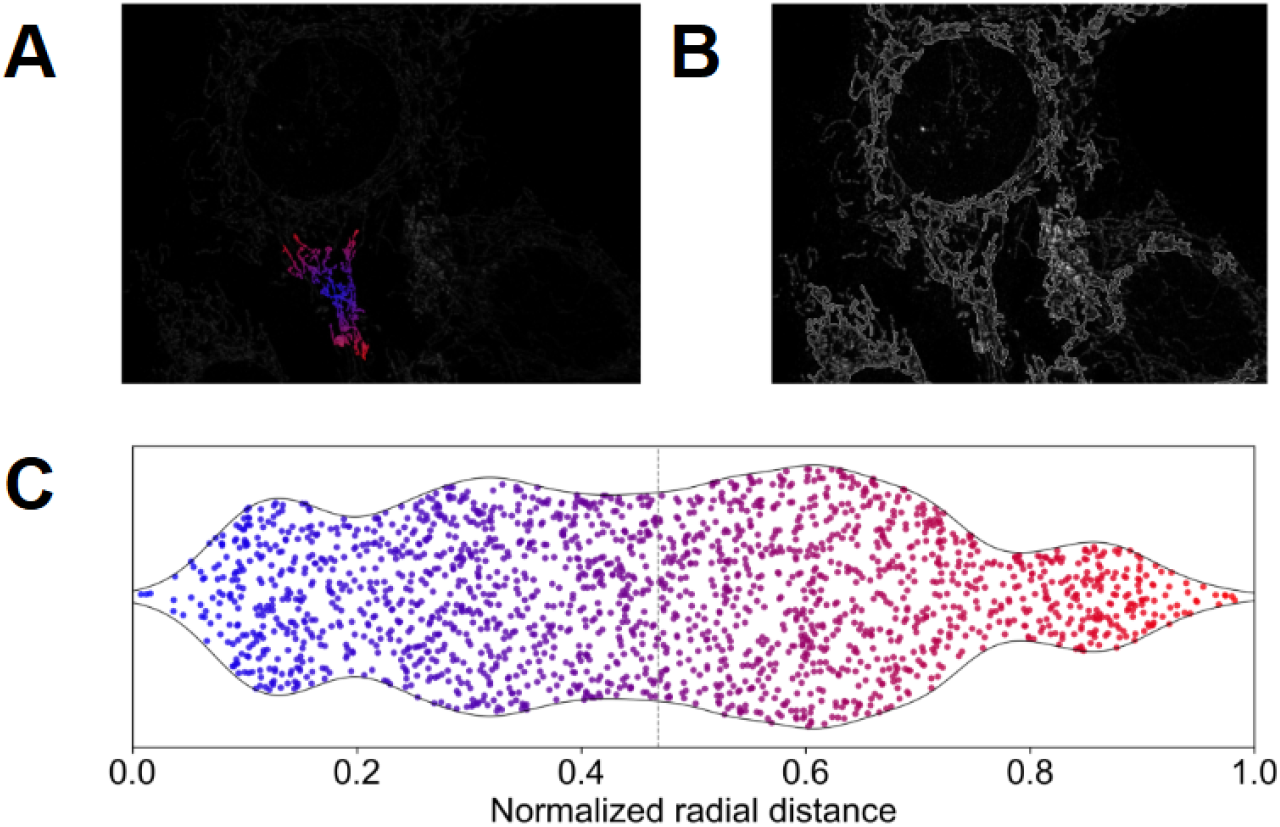
Mitochondrial localization in MEFs. **A)** Estimates of mitochondrial localization in a cell-cell contact network from center (blue) to periphery (red) from the anti-SDHA immunofluorescence (gray). **B)** Gray layer of cells and mitochondria detected in the image file. **C)** Violin plot with the fraction of mitochondrial pixels (dots) identified from center (0, in blue) to periphery (1, in red) for the highlighted region.

A third case study of the code consisted in the application to a *tif* file of HeLa cells expressing mito-YFP and stained for TOM20 from the public repository CLI under accession number 13731. We found that the code can automatically recognize mitochondria even when multiple mitochondria-targeted probes are mixed within the same channel (Figure 4).

**Figure 4.**
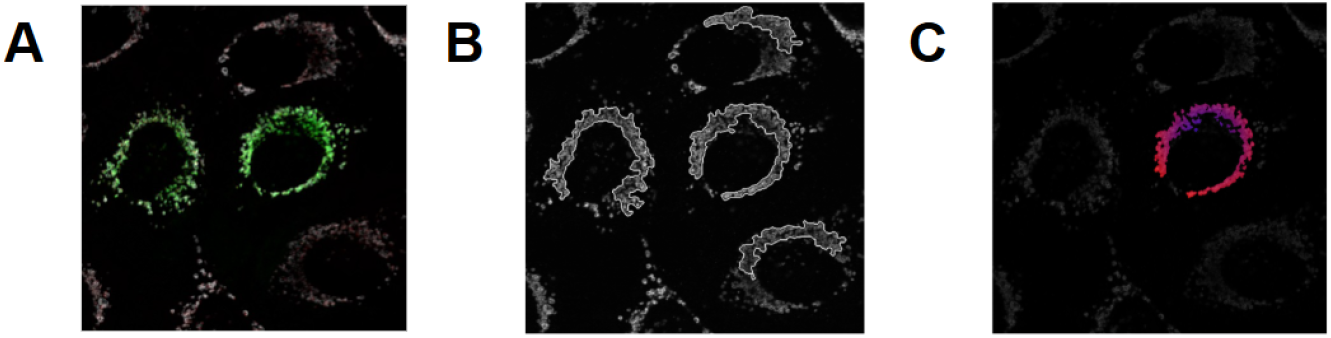
Mitochondrial localization in HeLa cells. **A)** Original *tif* figure showing HeLa cells colored by mito-YFP (green) and Tom20 (white) in the same channel. **B)** Gray layer of detected cells and mitochondria for both fluorophores in the single image of the file. The brightest parts of the cells are correctly detected. **C)** Per-cell estimates of mitochondrial localization from center (blue) to periphery (red).

Overall, the code performed well as a tool to assess mitochondrial distribution. Our tests indicated that the code infers cell boundaries exclusively from mitochondrial fluorescence, requiring only dense and broadly distributed mitochondrial labeling to approximate cellular area, as is typical of most cell types. It performs reliably with mitochondrial matrix–localized reporters, MitoTracker probes, or outer mitochondrial membrane markers that provide homogeneous cytoplasmic coverage. Importantly, touching cells may be segmented as a single cell in the absence of an independent nuclear or cytoplasmic reference channel. Consequently, the code is best suited for mitochondria-centric analyses rather than general-purpose cell segmentation.

## 4. Conclusions

We successfully generated a code capable of identifying and separating by peripheral or central location mitochondria within cells in images of different formats and with different mitochondrial stains. The code works well for images of spread, adherent cells with distributed mitochondrial networks. It does not require nuclear or membrane markers, nor a change of code for every individual analysis. We hope that automated and quantitative identification of mitochondrial cellular location can contribute towards better understanding of mitochondrial biology in health and disease.

## Acknowledgements

The authors thank Dr. Éverton Vogt for the assistance with MitoTracker microscopy assays, Prof. Alexandre Bruni Cardoso for microscopy access, as well as Dr. Julian Serna and Prof. Luca Scorrano (Univ. Padova) for donating immunofluorescence microscopies for analysis.

## Data Availability

The open-source code developed here was made fully available at https://github.com/cavalcantegc/mito_localization.git. Raw files are made available upon reasonable request.

## Author Contributions

G.C.C. – conceptualization, formal analysis, investigation, methodology, software, writing - original draft, writing - review & editing; A.J.K. – supervision, writing - review & editing. All authors read and approved the final manuscript.

## References

Benador IY, Veliova M, Mahdaviani K, Petcherski A, Wikstrom JD, Assali EA, Acín-Pérez R, Shum M, Oliveira MF, Cinti S, Sztalryd C, Barshop WD, Wohlschlegel JA, Corkey BE, Liesa M, Shirihai OS. Mitochondria Bound to Lipid Droplets Have Unique Bioenergetics, Composition, and Dynamics that Support Lipid Droplet Expansion. Cell Metab. 2018 Apr 3;27(4):869-885.e6. doi: 10.1016/j.cmet.2018.03.003. PMID: 29617645; PMCID: PMC5969538.

Glancy B, Kim Y, Katti P, Willingham TB. The Functional Impact of Mitochondrial Structure Across Subcellular Scales. Front Physiol. 2020 Nov 11;11:541040. doi: 10.3389/fphys.2020.541040. PMID: 33262702; PMCID: PMC7686514.

Haghighi M, McPhie D, Rohban M, Weisbart E, Logan DJ, Karhohs KW, Ewald JD, Haslum JF, Cimini BA, Singh S, Cohen BM, Carpenter AE. Identifying and targeting abnormal mitochondrial localization associated with psychoses. bioRxiv 2025.10.08.676630; doi: 10.1101/2025.10.08.676630

Jin SM, Lazarou M, Wang C, Kane LA, Narendra DP, Youle RJ. Mitochondrial membrane potential regulates PINK1 import and proteolytic destabilization by PARL. J Cell Biol. 2010 Nov 29;191(5):933–42. doi: 10.1083/jcb.201008084. PMID: 21115803; PMCID: PMC2995166.

Monzel AS, Enríquez JA, Picard M. Multifaceted mitochondria: moving mitochondrial science beyond function and dysfunction. Nat Metab. 2023 Apr;5(4):546–562. doi: 10.1038/s42255-023-00783-1. Epub 2023 Apr 26. PMID: 37100996; PMCID: PMC10427836.

Salazar MPR, Kolanukuduru S, Ramirez V, Lyu B, Manigault G, Sejourne G, Sesaki H, Yu G, Eroglu C. Mitochondrial fission controls astrocyte morphogenesis and organization in the cortex. J Cell Biol. 2025 Oct 6;224(10):e202410130. doi: 10.1083/jcb.202410130. PMID: 39484572; PMCID: PMC11527035.

Seager R, Lee L, Henley JM, Wilkinson KA. Mechanisms and roles of mitochondrial localisation and dynamics in neuronal function. Neuronal Signal. 2020 Jun 1;4(2):NS20200008. doi: 10.1042/NS20200008. PMID: 32714603; PMCID: PMC7373250.

Solomon T, Rajendran M, Rostovtseva T, Hool L. How cytoskeletal proteins regulate mitochondrial energetics in cell physiology and diseases. Philos Trans R Soc Lond B Biol Sci. 2022 Nov 21;377(1864):20210324. doi: 10.1098/rstb.2021.0324. Epub 2022 Oct 3. PMID: 36189806; PMCID: PMC9527905.

Vercesi AE, Castilho RF, Kowaltowski AJ, de Oliveira HCF, de Souza-Pinto NC, Figueira TR, Busanello ENB. Mitochondrial calcium transport and the redox nature of the calcium-induced membrane permeability transition. Free Radic Biol Med. 2018 Dec;129:1–24. doi: 10.1016/j.freeradbiomed.2018.08.034. Epub 2018 Aug 31. PMID: 30172747.

